# Binding Structures, Mechanical Properties, and Effects on Cellular Behaviors of Extracellular Matrix Proteins on Biomembranes

**DOI:** 10.64898/2026.04.03.716427

**Authors:** Veronica Ivanovskaya, Jessica Ruffing, Minh D. Phan

## Abstract

Extracellular matrix (ECM) proteins assemble to form a heterogeneous connective scaffold that supports cells. Physical interactions between cells and the matrix regulate cellular behaviors and influence subsequent tissue construction. However, there is a lack of fundamental understanding regarding the contributions of individual native ECM proteins to the matrix. This gap arises from the need for nanoscopic characterization, which operates on a much smaller length scale than typical assessments in cell and tissue cultures, as well as in tissue reconstruction and clinical implantation. This study aims to systematically investigate how individual ECM proteins affect lipid membranes structurally and mechanically, and how these influences regulate cell migration. Results from Langmuir isotherm analysis, X-ray reflectivity measurements, and cell scratch assays demonstrate that strong collagen adsorption on the membrane surface disrupts lipid packing. However, its rigid network provides a sturdy scaffold for cell adhesion, thereby enhancing cell attachment and promoting cell migration. In contrast, elastin has a minimal structural or mechanical impact on the membrane during both adsorption and compression, but it benefits cells by facilitating migration and reducing the risk of infection. Fibronectin, on the other hand, exhibits complex mechanical responses to compression, characterized by significant structural rearrangements that occur during adsorption. This strong interaction with the membrane can result in excessively high adhesion forces, ultimately limiting cell motility. These findings lay the foundation for the design of artificial scaffolds that can manipulate cellular responses, a critical step toward advancing regenerative medicine and tissue engineering.

**Significance:** Fabricating extracellular matrix (ECM) scaffolds from cells offers advantages over traditional approaches, such as decellularized tissues, which face donor limitations, and artificial scaffolds, which may hinder cellular communication. However, the slow harvesting process of cell-derived ECM has limited its clinical applications. This research is part of a larger mission to engineer ECM prescaffolds on lipid carriers tailored to cell requirements, enhancing ECM production and regulating cell behavior. The first step involves systematically analyzing the structural and mechanical effects of ECM on lipid membranes and how these effects regulate cellular behavior. This work confirms distinct characteristics of ECM proteins, advancing fundamental understanding of cell-matrix interactions and paving the way for scaffold engineering.

## Introduction

Regenerative medicine focuses on restoring the functionality of lost or damaged tissues by helping the body heal. This field includes gene therapy, cell therapy, specialized drug delivery, and tissue engineering, which use various natural and synthetic materials. Tissue engineering often employs scaffolds, platforms for tissue formation, that can be implanted in the body. Effective scaffolds must be biocompatible, biodegradable, highly porous, and possess adequate mechanical properties to support cell adhesion and growth [1]. Currently, scaffold options include premade porous scaffolds, hydrogel scaffolds, decellularized extracellular matrices and cell-derived extracellular matrices [2]. Pre-made porous scaffolds, while diverse in material, often use artificial polymers that possibly hinder cellular communication [3, 4]. Decellularized extracellular matrix scaffolds provide the most native-like environment, but require organ donors, making them challenging to produce. Cell-derived extracellular matrix sheets do not depend on organ donors but have limited applications due to their long production timelines. Ultimately, while various scaffolds exist, extracellular matrix sheets have greater potential to replicate the native extracellular matrix required for effective cell growth once optimized matrix secretion efficiency is achieved.

The extracellular matrix (ECM) is a network of proteins, glycosaminoglycans (GAGs), and proteoglycans synthesized within cells and released through exocytosis [5, 6] primarily composed of collagen, elastin, fibronectin, and laminin, with variation in composition among various cell and tissue types [7]. Often described as the glue that connects cells, ECM plays a vital role in cell differentiation, adhesion, and migration, while also regulating the mechanical properties of a cell’s environment [8]. The outermost layer of the ECM, the interstitial matrix, contains structural proteins such as collagen types I to III, elastin, and additional proteoglycans [7, 9]. These collagens are hydrophobic, stiff, rod-shaped proteins that resist mechanical stress, accounting for about 30% of the total protein mass of the body [7]. Collagen I is essential for providing mechanical strength and stability to the ECM [10], and is deposited at higher concentrations in tissues that need stiffness, such as cartilage and bone. Cells rely on mechanical stimuli from the ECM through mechanotransduction to sense their environment, which affects their migration and adhesion [11]. Elastin, another hydrophobic structural protein found in the interstitial matrix, is characterized as a randomly coiled structure with some covalent crosslinking of hydrophilic lysine residues, giving it amphiphilic properties. As an elastic natural polymer, it provides low stiffness, high extensibility, and efficient storage of elastic energy in the ECM [12].

Elastin is abundant in soft, elastic tissues, including the vascular system, lungs, skin, and bladder. It contributes to biological processes such as migration signaling, cell proliferation, adhesion, and growth [13]. Studies suggest that elastin can improve the fibrillogenesis of collagen I [14] and that fibronectin accelerates the deposition of these two proteins [15, 16]. A large dimeric glycoprotein (230-270 kDa), fibronectin serves as a transition molecule, meaning it can be found in both the interstitial matrix and the basement membrane, the lowest layer of the ECM, which consists mainly of laminins, collagen IV, nidogens, and heparan sulfate proteoglycans [17]. Fibronectin, possessing multiple binding domains connected by cysteine disulfide bonds [18], acts as a regulatory protein, binding to growth factors, other proteins, and itself in response to cytoskeletal tensile forces [19]. The protein naturally prefers a folded state when suspended in solution, but will unfold upon electrostatic interactions [20, 21] or under mechanical forces [22, 23]. Fibronectin organizes the interstitial matrix through signaling, adhesion, migration, and morphogenesis, and regulates mechanical properties similarly to collagen. Its assembly is influenced by the basement membrane, which anchors ECM proteins with transmembrane proteins that facilitate mechanical and chemical signaling with cells [24–26].

These native polymers are commonly incorporated into hydrogels and other types of scaffolds to mimic the ECM. However, the composition of the extracellular environment evolves over time and is much more complex than that of current scaffolds, making the development of accurate platforms challenging [7, 27]. ECM secreted by confluent cells possesses the desired level of sophistication [27]. However, producing it is a slow process that cannot be optimized without methods to manipulate the resulting matrix and its modifications. The restructuring of the ECM occurs in response to environmental stimuli, involving the synthesis or degradation of various molecules to alter its mechanical and/or chemical properties. Despite significant variations in the mechanical properties and structures of proteins within the ECM, predicting how restructuring will occur is difficult due to the overlapping functions of these proteins.

By understanding the mechanical contributions of individual proteins to the ECM, we can create optimized scaffolds. This can be achieved by supplementing the ECM with molecular components and manipulating modifications alongside cellular responses. With this knowledge, growing cells can be supplemented with lipid nanoparticles containing specific prescaffold native ECM proteins, which would serve as instructions to direct the cells’ ECM secretion. The most prevalent ECM proteins – collagen, elastin, and fibronectin – exhibit unique mechanical properties, making them suitable for developing diverse instructions. In this study, our objective is to assess the structures and mechanical effects of individual ECM proteins on lipid membranes using Langmuir isotherms and X-ray reflectivity, and to examine these influences on cellular migration through scratch-assay experiments. The results demonstrate the distinct protein adsorption, structural, and mechanical behaviors of individual ECM proteins, as well as the advantages of formulating ECM prescaffolds on lipid membranes, rather than free, native forms of these proteins in solution, when examining cell proliferation, migration, and susceptibility to infection.

## Materials and Methods

### Lipid Monolayer Preparation

14:0 1,2-dimyristoyl-sn-glycero-3-phosphocholine (DMPC) and 14:0 1,2-dimyristoyl-sn-glycero-3-phospho-L-serine (DMPS) lipids obtained from Avanti Polar Lipids (Alabaster, AL) were used to create a negatively charged lipid monolayer for the characterization of lipid-protein interactions. DMPC and DMPS were mixed together at a 4:1 mole ratio and dissolved in a mixture of chloroform and methanol at a 5:1 volume ratio to achieve a 1 mg/mL concentration solution that would then be used with elastin from bovine neck ligament purchased from Sigma Aldrich (St. Louis, MO) and human plasma fibronectin from Thermo Fisher Scientific (Carlsbad, CA). This mixture was selected to create a negatively charged surface, facilitating protein adsorption and the unfolding of ECM proteins onto the lipid platform [21]. In order to work with rat type I collagen from Sigma Aldrich (St. Louis, MO), plant cholesterol was also obtained from Avanti Polar Lipids (Alabaster, AL) and added to a DMPC:DMPS mixture at a 25 mol% to promote protein association, based on our previous findings [28]. Lipid monolayers were formed using an asymmetric film compression 32.8 × 9.7 *cm*^2^ Langmuir trough system (KSV NIMA 302 M; Biolin Scientific, UK) that was thoroughly cleaned prior to each use by wiping with non-sterile cleanroom wipers soaked in chloroform or ethanol (for Teflon surfaces), and rinsed with MilliQ water. Any remaining water on the trough or barriers was aspirated. The value of surface pressure (Π), calculated from the difference between the bare buffer surface tension and the film-covered surface tension, was monitored by a Wilhelmy plate that hung from a high precision microbalance as given by,

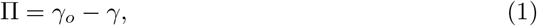

where *γ*_*o*_ is the surface tension of the air-water interface and *γ* is the surface tension in the presence of a monolayer film. The platinum Wilhelmy plate was blowtorched to remove any residual solvent, protein, or lipid. The trough was then placed in its plexiglass box with two Kapton windows on an active antivibration table within an X-ray reflectometer (D8 Advance; Bruker, Germany). The trough was filled with a filtered 0.05 M Tris-HCl buffer at pH 7.2, ensuring the liquid level rose above the edges to form a positive meniscus and allowing full contact with the barriers and the Wilhelmy plate. The working surface was cleaned by first closing the barriers, then using gentle aspiration to remove surface contaminants until the compression and expansion of the barriers maintained a constant pressure. To form the monolayer, the lipid mixture was tapped onto the buffer surface in small drops using a Hamilton syringe. The system was given ten minutes for the organic solvents to evaporate and for the monolayers to stabilize before the Π-A isotherms were recorded at a constant compression rate of 2 mm/min.

### Mechanical Properties

In order to assess two-dimensional elastic packing interactions within the lipid monolayer, monolayer compressibility *C*_*s*_ was calculated using,

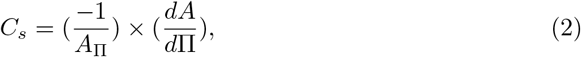

where Π is the surface pressure, and *A*_Π_ is the molecular area at the calculated Π. Data was collected at 1-second sample intervals. The resulting Langmuir isotherm data were then extracted prior to film collapse and averaged across repeated experiments. Further analysis uses the reciprocal of monolayer compressibility, the elastic modulus of the area compressibility 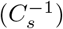, allowing for quantitative comparison with bilayer lipid systems. The higher modulus indicates the film’s greater resistance to area compression, providing insight into the interfacial elasticity arising from lipid-lipid and lipid-protein interactions [28–31]. This method is typically used for amphiphilic molecules, including surfactants, lipids, and polymers, but it can also be applied to less amphiphilic molecules, such as proteins, as long as these molecules are attached to a free-standing platform on the water’s surface. Data points nearest to half- and whole-integer values were selected, isolated, and smoothed using LOESS, a nonparametric method that does not require prior trend assumptions, revealing unique trends.

### ECM Protein Adsorption Assays and Compression Isotherms

Each of the three proteins was diluted in filtered Tris-HCl buffer and adjusted to various concentrations and pH values. The protein solutions were prepared with final concentrations of 2.9 *µ*g/mL elastin (pH 8.8), 5.69 *µ*g/mL collagen supplemented with 2–3 drops of acetic acid as per the manufacturer’s recommendation, and 17.6 *µ*g/mL fibronectin for a 1:5 protein to lipid ratio. These solutions were vortexed prior to use and deposited after the lipid monolayer had been compressed to 12.5 mN/m. This was held constant for 15 minutes to equilibrate the monolayer structure, then switched to fixed surface area (stopped the barriers) while monitoring surface pressure changes.

Without disturbing the surface or increasing the surface pressure by more than 0.15 mN/m, 1 mL of the individual protein was injected beneath the film by carefully sliding the needle beneath the barriers. The proteins were allowed to adsorb, and changes in surface pressure corresponding to protein adsorption were recorded over 24-72 hours. To minimize evaporation during longer studies, a small dish containing buffer-damped Kimwipes was kept within the plexibox to maintain humidity. The evaporation level was confirmed by X-ray reflectivity, with less than 50 *µ*m change in the water surface position over 65 hours, resulting in less than 0.5 mN/m change in the surface pressure. For the compression isotherms, after protein deposition, the barriers were set to compress at 0.5 mm/min, and the Langmuir isotherm profiles were recorded until lipid film collapse.

### X-ray Reflectometry

X-ray reflectometry (XRR) measurements were performed at a wavelength of *λ* = 1.54 Å from a Cu K*α* source, using a vertical goniometer and a horizontal sample geometry, on an undisturbed liquid surface. A parallel incident beam was produced by a Göbel mirror and used two slits to define the size of the incident beam and reduce its vertical divergence. Intensity was measured as a function of momentum transfer (*q*_*z*_),

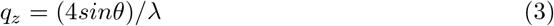

where *λ* and *θ* are the wavelength and incident angle, respectively. Reflectivity was measured over a range of *q*_*z*_ values, allowing determination of thickness and density changes within the protein-lipid layers. X-ray scattering length density (SLD) is a product of *r*_*e*_*× ρ*, where *r*_*e*_ is the classical electron radius (2.814 *×*10^*−*5^ Å) and *ρ* is the electron density. The X-ray SLD profiles perpendicular to the reflecting interface were acquired from reflectivity data curve fits. In this work, XRR scans were performed on the bare lipid membrane at 12.5 mN/m before and after protein deposition, and after a minimum of 15 hours when the adsorption reached equilibrium. The protein-bound lipid films were then further compressed to pressures of 20 mN/m and 30 mN/m, and XRR was measured accordingly. The data were then analyzed using amoeba and chi-squared fittings in LSFit and DREAM fitting in Refl1D to obtain error ranges. Analysis was performed using a two-layer model, which was later replaced with a more sophisticated three-layer model once it became unrepresentative of the observed protein adsorption system.

### Scratch Assays

HDFn cells obtained from American Type Culture Collection (ATCC) were cultured in accordance with the vendor’s recommendations with Fibroblast Basal Media (FBM)(Manassas, VA) from ATCC and supplemented with a Serum Free Fibroblast Growth Kit (Manassas, VA) from ATCC. Passaging was performed every 3 days to provide cells with sufficient nutrients and space to continue proliferating. HDFn cells were passaged four times and seeded into 12-well culture plates at a density of 2,500–5,000 cells/cm^2^, with 2 mL of growth medium added per well. Each well had a horizontal line drawn to reference during scratching. Lipid monolayers were prepared in previously explained methods with 14:0 1,2-dimyristoyl-sn-glycero-3-phosphocholine (DMPC) and 14:0 1,2-dimyristoyl-sn-glycero-3-phospho-L-serine (DMPS), and plant cholesterol obtained from Avanti Polar Lipids (Alabaster, AL). Elastin from bovine neck ligament purchased from Sigma Aldrich (St. Louis, MO), human plasma fibronectin from Thermo Fisher Scientific (Carlsbad, CA), and rat type I collagen from Sigma Aldrich (St. Louis, MO) were individually adsorbed onto the lipid membranes as previously explained to create an ECM scaffold. Lipid membranes act as carriers of native ECM proteins to cells [32]. Varying experimental conditions were assigned to each well. The protein–lipid mixture was added to achieve a final concentration of 20% (400 µL per well). Conditions were diluted to achieve 10 *µ*g of added protein, with and without vesicles. Cells were grown to 90-100% confluence in media without supplementation from a growth kit. Once grown, a vertical scratch was made using a 200 *µ*l pipette tip. Images at 10x magnification were taken from the zeroth to the fifty-fourth hour, with the first hour at 3-hour increments and the remaining hours at 6-hour increments. Migration analysis was performed in ImageJ by outlining each scratch with the polygon tool and measuring the scratch area in *µ*m using the measure tool (Image processing using ImageJ is described in S1, Supplemental Information).

MATLAB code was developed to extract pixels in the RGB color model with values [195, 255, 0] with a tolerance of 0.3, and to produce a grayscale image or a mask of pixels in this range. The pixels in the mask were compared to the total number of pixels to yield the amount of infection within the cells (Image processing using MATLAB is described in S2, and scratch assay images at selective hours are provided in S3, Supplemental Information).

## Results

### Two-Dimensional Lipid Film Characterization

Characterizing the lipid monolayers establishes a reference point for evaluating how ECM proteins influence lipid packing and the resulting mechanical behavior. Langmuir isotherms provide a two-dimensional description of the lipid film, capturing not only its mechanical response but also the thermodynamics associated with phase transitions. Upon compression, the monolayer progresses from a highly disordered gaseous phase to more densely packed liquid and solid phases. The Π-*A* isotherms in Figure S4 (Supplemental Information) illustrate how DMPS and cholesterol interact with DMPC. Moreover, the Langmuir setup (Fig. 1A) enables analysis of the thermodynamics of mixing in multicomponent systems via their packing characteristics, the mechanical and rheological properties of the films through their resistance to compression, and the use of a precisely controlled film surface area or packing density for adsorption experiments and surface scattering measurements.

**Figure 1.**
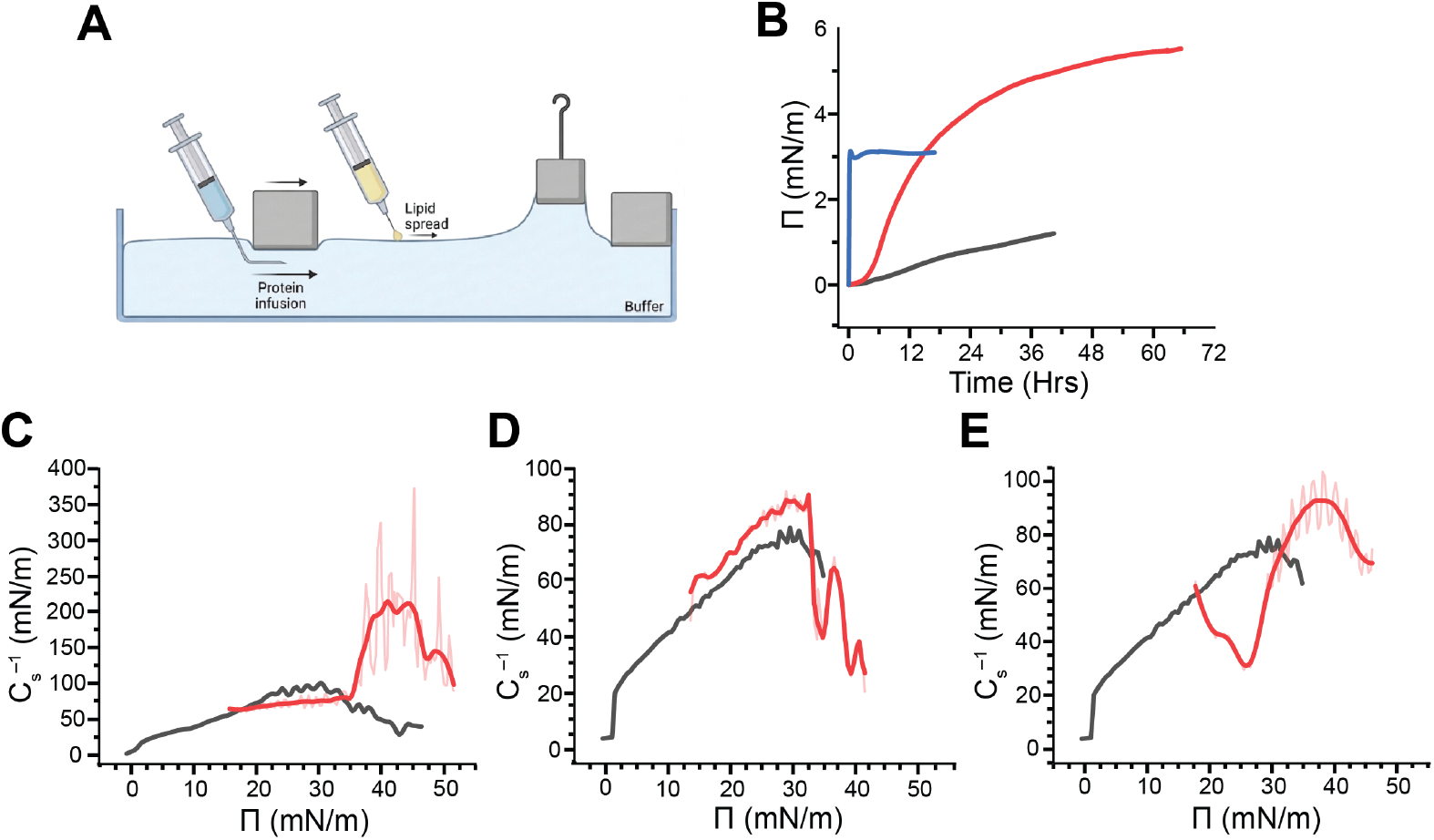
(A) Langmuir apparatus schematic. (B) Adsorption (Π vs time) of individual proteins to their respective lipid films. The blue curve represents collagen, the yellow represents elastin, and the red represents fibronectin. Mechanical response of individual lipid-adsorbed proteins (C) collagen, (D) elastin, and (E) fibronectin systems to compressive forces. The curves were smoothed using LOESS to track trends without losing data accuracy.

### Protein Adsorption Assays

Adsorption assays provide insight into protein-lipid interactions, allowing observation of parameters such as protein binding affinity, lipid film adsorption capacity, and adsorption kinetics. Figure 1B shows the adsorption assays of each ECM protein as a function of surface pressure change that took place after injection. Collagen adsorption occurs instantaneously, increasing surface pressure by 3.1 mN/m, followed by a sharp decline to 0.4 mN/m. The adsorption curve gradually rises back to its maximum surface pressure, then mildly decreases again, forming a broad peak over 12.5 hours and plateauing thereafter. Elastin exhibits steady adsorption for 40 hours, with a slow increase in surface pressure by only 1.2 mN/m. Fibronectin’s adsorption curve followed a sigmoidal trend, reaching saturation at approximately 65 hours with a total surface pressure change of 5.5 mN/m. Note that the adsorption kinetics obtained from the Langmuir apparatus may take longer than similar experiments where both lipid vesicles and proteins are dispersed in solution [21], because it not only depends on lipid-protein interactions but also on the amphiphilicity of proteins that drives them to the water surface.

### Protein Influence on Lipid Packing and Mechanical Properties

Interfacial elasticity is commonly used to characterize lipids, but can also be applied to other amphiphilic molecules forming a free-standing film or adhering to a platform at the water’s surface. The versatility of this method enables the mechanical properties of a wide range of materials to be determined, including ECM proteins, whose mechanical properties are vital for regulating cellular mechanical behavior. As shown in Figure 1C, collagen’s deposit slowly increases elastic modulus beginning at 64.4 mN/m, and rising to 81.9 mN/m at a surface pressure of 35 mN/m. During compression, the addition of collagen lowers the initial elastic modulus of the collagen-coated membrane and slows its increase as surface pressure increases, as compared to the pure membrane modulus; however, beyond 35 mN/m, the system significantly stiffens, exhibiting a 160% increase in elastic modulus and reaching a peak value of 255 mN/m at a surface pressure of 42 mN/m. The latter behavior is expected to be dominated by collagen’s intrinsic rigidity. In contrast, the mere introduction of elastin does not change the elastic modulus of the lipid system significantly (Fig. 1D). Once compression begins, the overall system shows an increase in elastic modulus to 10 mN/m, but at a surface pressure of 30 mN/m, it triggers an amplified collapse and a decrease in stiffness, a characteristic expected of elastin. The compression of fibronectin, similar to that of collagen, softens the lipid film during the 17.8-25.8 mN/m surface pressure range as indicated by the two local minima and elastic modulus of 30 mN/m (Fig. 1E). The fibronectin-coated membrane stiffens sharply above 30 mN/m, delaying the collapsing surface pressure by 7.4 mN/m. Both elastin and fibronectin increase the elastic modulus by only 14% from just a pure lipid film with maximum moduli of 91 mN/m.

### X-ray Reflectometry of Individual Protein Systems

X-ray reflectometry (XRR) complements the characterization of the protein–lipid system along the third dimension, the z-axis, by recording the intensity of X-rays reflected from individual layers within a free-standing film. Due to the specific atomic composition of lipid molecules, XRR can differentiate between headgroup and tail regions and can detect structural rearrangements following protein adsorption. XRR data were acquired before protein addition, immediately after protein injection, and once protein adsorption had reached equilibrium. Subsequently, the protein-covered lipid films were compressed to surface pressures of 20 and 30 mN/m, and XRR was recorded at each pressure. Two-layer model X-ray scattering length density (SLD) profiles were evaluated against those of the pristine lipid membrane to determine how protein binding alters membrane architecture. When the two-layer fit no longer adequately represented the measured profiles, a three-layer model was employed instead. In the X-ray SLD depth profiles, z = 0 is defined at the midpoint of the lipid headgroup region; negative z values correspond to the protein and subphase side, while positive z values extend toward the lipid tails and air.

At the time of deposition, the collagen-coated membrane (red profile, Fig. 2A) exhibits an oscillation fringe shifted toward lower *q*_*z*_ values and a higher reflectivity compared to the bare membrane (black, Fig. 2A). This corresponds to an increase in both thickness and material density in the lipid headgroup region (red, Fig. 2B), consistent with the adsorption assay results. After allowing additional time for adsorption and equilibration (blue, Fig. 2A), the oscillation fringe in the XRR curve shifts slightly further toward lower *q*_*z*_, indicating a further increase in overall sample thickness. The corresponding fit suggests that collagen molecules migrate into the region between the head and tail groups, which is reflected by an additional increase in thickness combined with a decrease in the material density of the headgroup region, as well as a reduced tail-region thickness due to packing disruptions (blue, Fig. 2B). Once compression begins (green and purple, Fig. 2A), the oscillation fringes move further to lower *q*_*z*_, and the reflectivity diminishes much more rapidly than in the earlier stages, indicating increased interfacial roughness and layer thickness, along with a potential decrease in packing density within the layers. The fitted models corroborate this by showing reduced X-ray SLDs and a pronounced rise in interfacial roughness, which together imply disrupted packing in both the headgroup and tail regions (green and purple, Fig. 2B).

**Figure 2.**
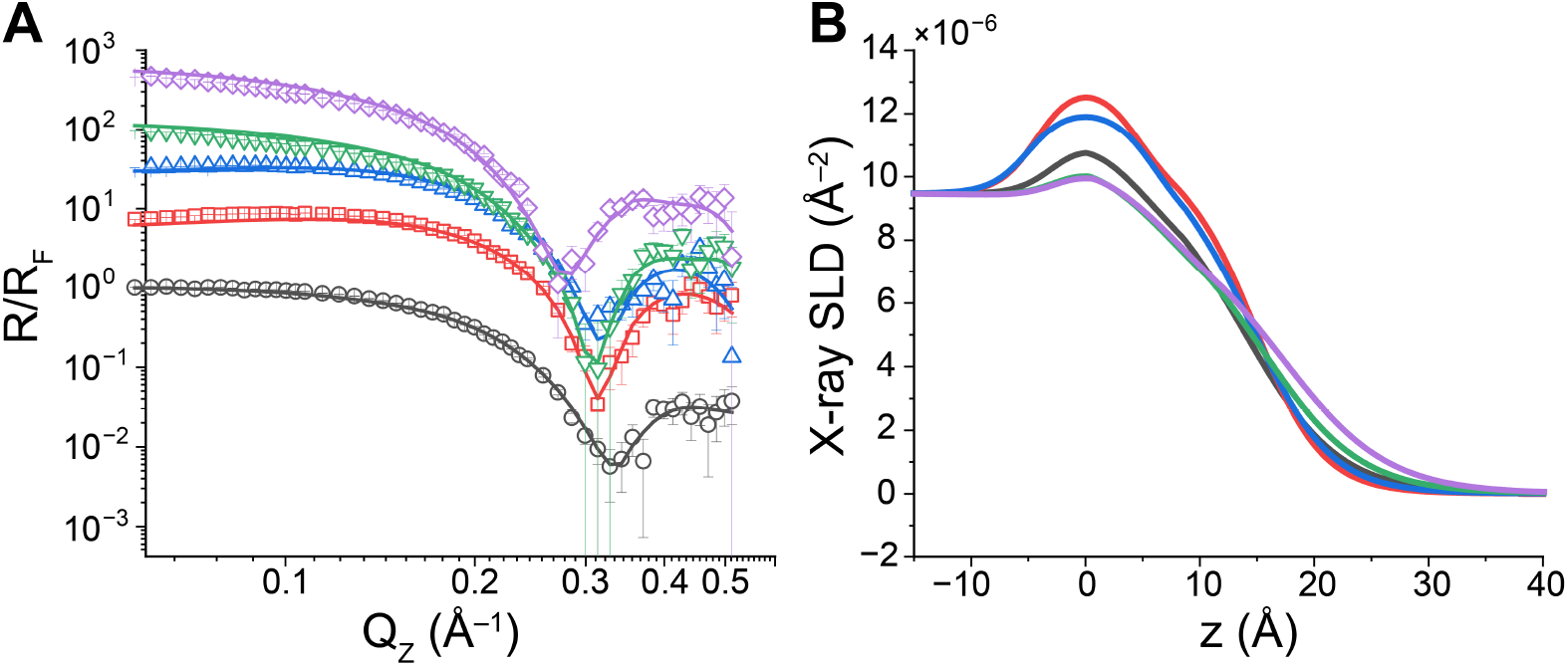
(A) X-ray reflectivity data for the lipid membranes before (black) and immediately following (red) collagen deposit, after adsorption (blue), and further compressing at 20 mN/m (green) and 30 mN/m (purple) surface pressures. The lines were generated using an iterative procedure and are the best fits, resulting in (B) the corresponding X-ray scattering length density (SLD) profiles.

**Figure 3.**
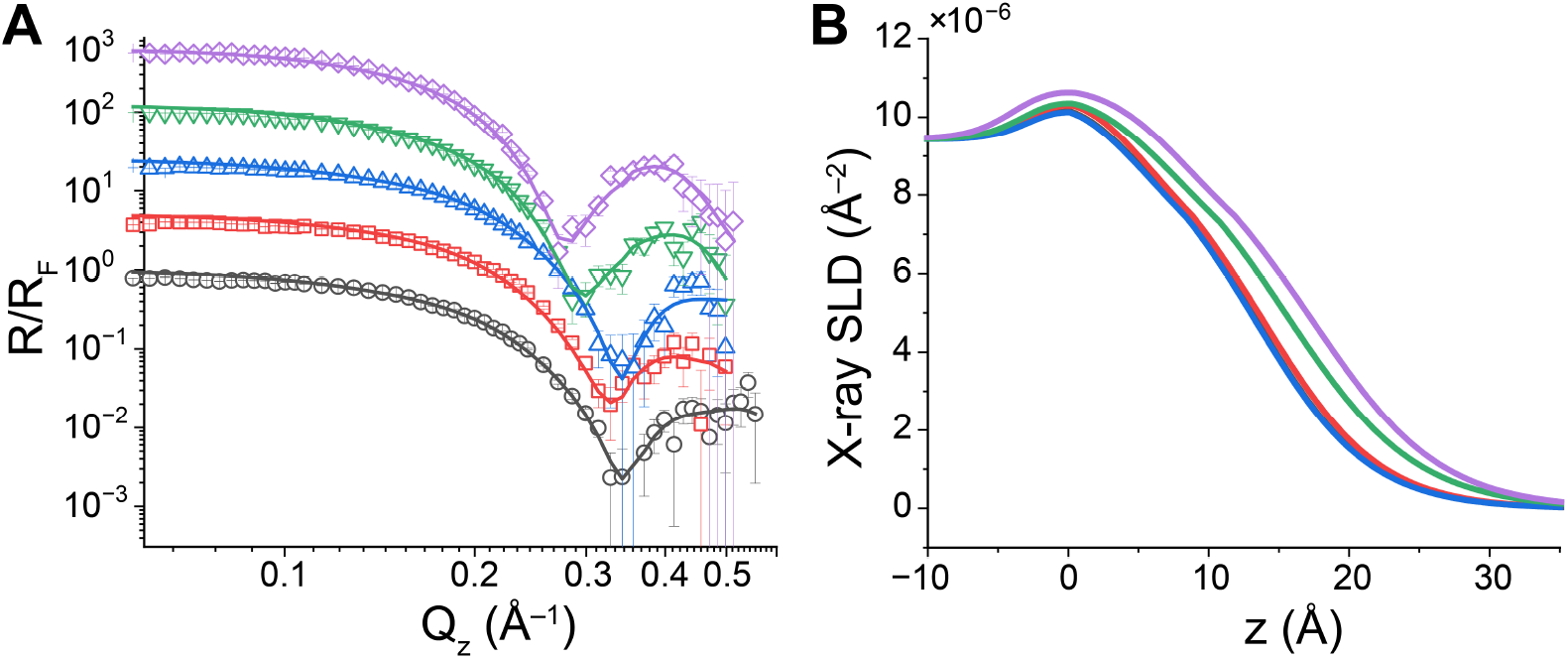
(A) X-ray reflectivity before (black), immediately following elastin deposit (red), after adsorption (blue), and further compressing at 20 mN/m (green) and 30 mN/m (purple) surface pressures. The lines were generated using an iterative procedure and are the best fits, resulting in (B) the corresponding X-ray scattering length density (SLD) profiles.

Upon elastin deposition (red, 3A), the XRR profile shifts slightly toward lower *q*_*z*_ values without notable changes in reflectivity, indicating an increase in the overall sample thickness. The corresponding fit (red, 3B) verifies that the headgroup thickness increases, while the lipid head–tail interfacial roughness decreases slightly, consistent with non-disruptive interactions between elastin molecules and the headgroup region, thereby promoting packing uniformity. This non-disruptive interaction is further evident at the adsorption equilibrium, where both the XRR profile and the fitted model are nearly indistinguishable from those of the bare membrane. Under compression, the XRR profile shifts further toward lower *q*_*z*_, reflecting a further increase in total thickness (green and purple, 3A). The corresponding fitted models resolve more detailed structural changes: the headgroup thickness and interfacial roughnesses increase, whereas the tail thickness remains constant, implying that elastin is incorporated into the lipid head region and extends to the head–tail interface (green, 3B). By the end of compression, both head and tail thicknesses increase, while the two interfacial roughnesses decrease, pointing to improved layer-by-layer packing uniformity. The non-disruptive behavior of elastin is further underscored by the X-ray SLD values of the head and tail regions, which remain unchanged throughout compression.

Similar to collagen, fibronectin interacts strongly with the lipid monolayer and drives pronounced alterations in membrane structure. Immediately after injection, the XRR profile (red, Fig. 4A) exhibits nearly the same fringe position as the bare membrane but with a modest increase in reflectivity, consistent with a higher material density. The corresponding fit (red, Fig. 4B) indicates that the X-ray SLD of both the head and tail regions increases, with a slight thickening of the headgroup layer and a small reduction in the tail thickness, while the overall thickness and roughness of the film remain essentially unchanged. At adsorption equilibrium, the XRR profile (blue, Fig. 4A) displays a further, though small, increase in reflectivity intensity together with the loss of the oscillatory fringe. This suggests a rise in average material density and either a diminished SLD contrast between layers or an increase in interfacial roughness on the scale of the layer thickness. The fit (blue, Fig. 4B) supports the latter interpretation, showing that FN is located between the head and tail regions and induces substantial disorder in the tail layer.

**Figure 4.**
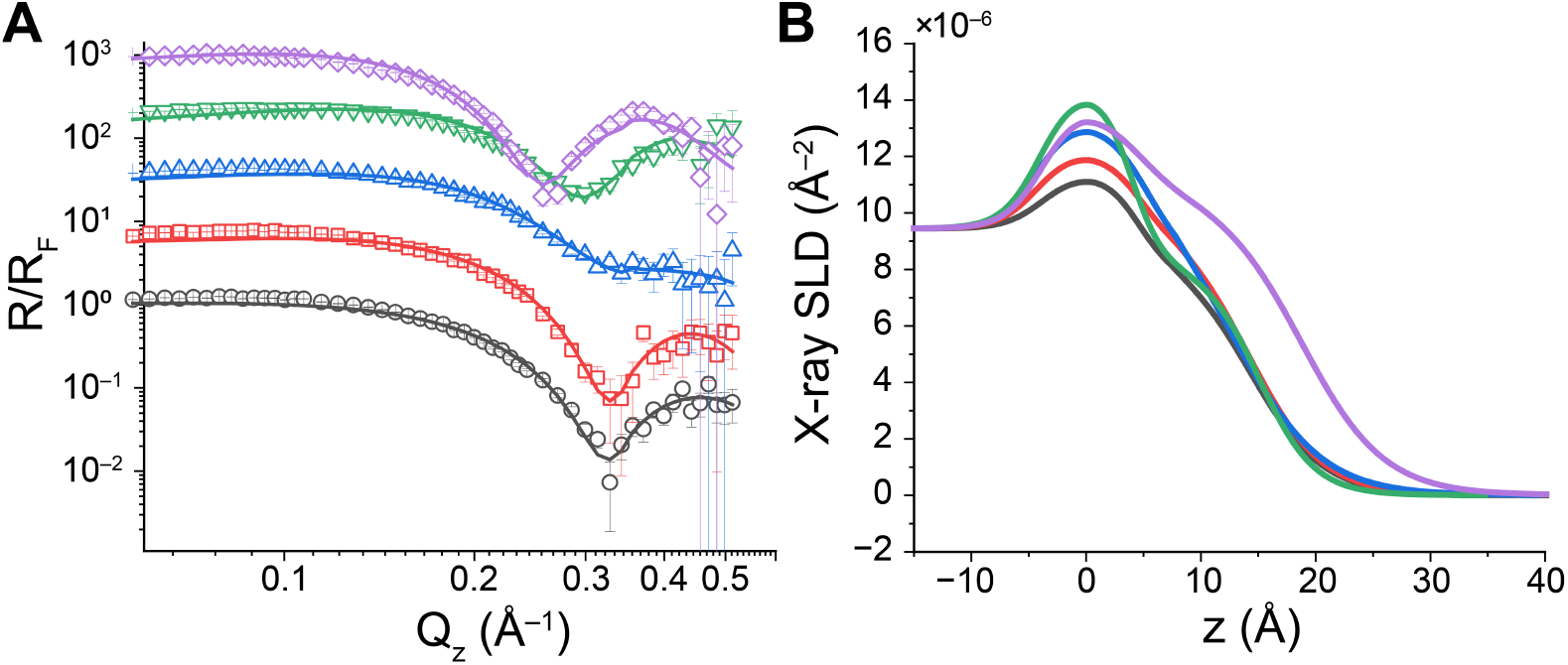
(A) X-ray reflectivity before (black), immediately following fibronectin deposit (red), adsorption equilibrium (blue), and further compressing at 20 mN/m (green) and 30 mN/m (purple) surface pressures). The lines were generated using an iterative procedure and are the best fits, resulting in (B) the corresponding X-ray scattering length density (SLD) profiles.

Upon compression (green, Fig. 4A), the reflectivity intensity rises again, and the oscillatory fringe reappears, implying a further increase in material density. In addition, the adsorbed FN molecules reorganize, likely enhancing the SLD contrast between adjacent layers. The fitted model (green, Fig. 4B) indicates a partial recovery of the membrane architecture from the disruption observed at adsorption equilibrium. The headgroup SLD increases, while both its thickness and interfacial roughness decrease, consistent with FN molecules occupying voids in the headgroup region and thereby increasing its density. Conversely, the tail SLD decreases to a value close to that of the pristine membrane, while its thickness increases markedly, suggesting that FN redistributes toward the headgroup region and allows the lipid tails to repack more normally under compression. Additional compression causes a further shift of the fringe toward lower *q*_*z*_ values (purple, Fig. 4A), corresponding to an increase in both tail thickness and SLD, as reflected in the fitted model (purple, Fig. 4B).

### Cellular Migration and Infection Assessment by Scratch Assays

After characterizing the structural and mechanical properties of the deposited ECM proteins, we used the same formulations in liposome systems to examine their influence on cellular behavior. We conducted *in vitro* experiments with human neonatal dermal fibroblasts (HDFn), which are key contributors to tissue repair. Scratch assays are commonly employed in wound-healing research to monitor cell migration under various treatment conditions. Following tissue injury, secreted cytokines recruit fibroblasts from surrounding regions [21, 33]. During this process, fibroblasts become activated and move toward the damaged area, in parallel with the accumulation of ECM proteins [34, 35]. In our experiments, we applied ECM protein-coated liposomes and free soluble ECM proteins to the wounded area as treatments. Figure 5A presents a schematic of the scratch, which is partitioned into two halves within a single well. Figure 5B and C depict the temporal progression of wound closure by cell migration through representative images at 0 and 54 hours, and also show the emergence of bacterial contamination (bright neon dots) observed at the conclusion of the scratch assay (Fig. 5C). The two control samples consisted of: 1) Tris-HCl buffer only without liposomes and ECM proteins, and 2) a liposome solution without ECM proteins (Fig. S3, Supplemental Information).

**Figure 5.**
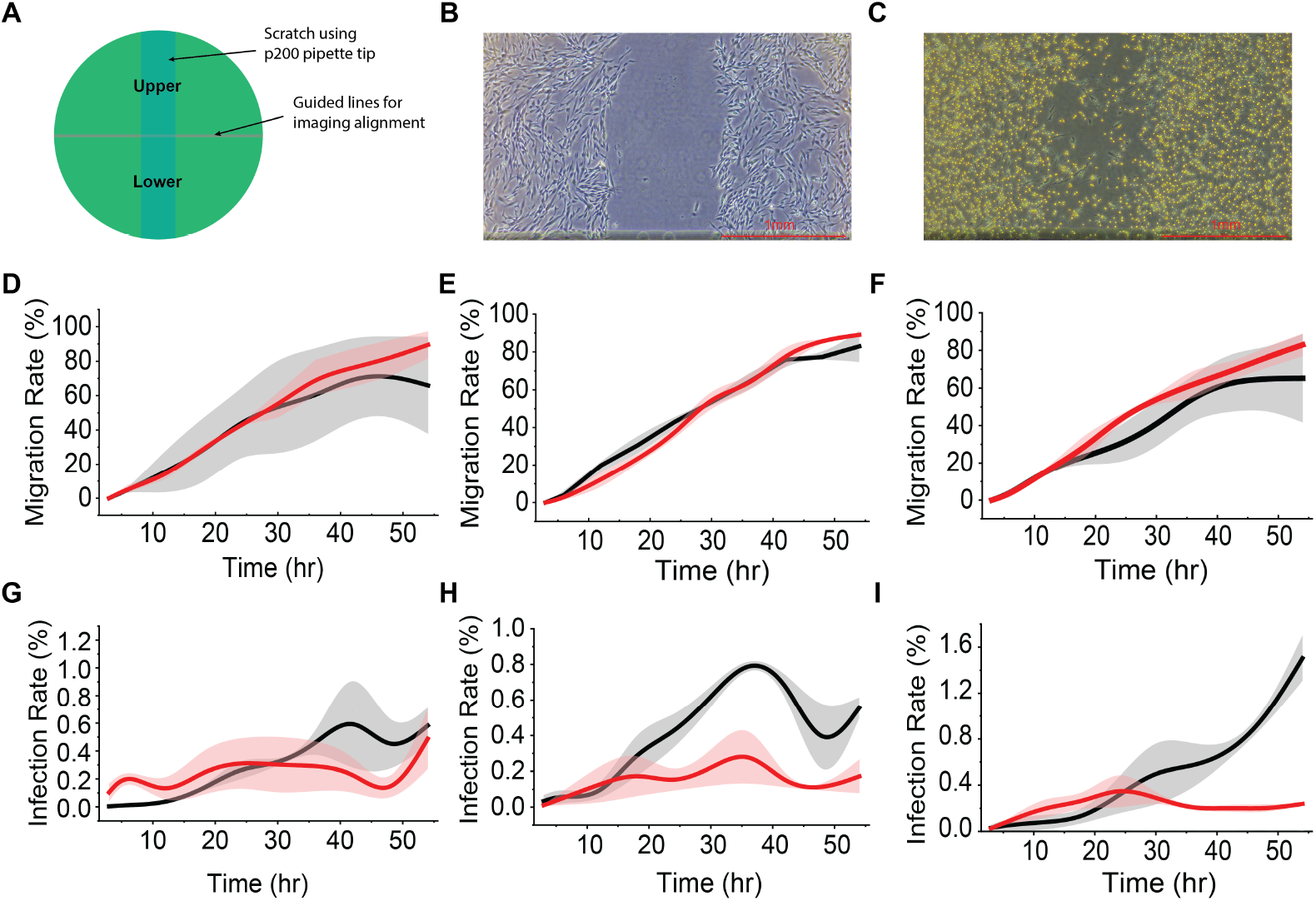
Evaluation of cell migration and infection rates in collagen, elastin, and fibronectin formulations. The scratch was performed at hour 0 and monitored for 54 hours. Each well plate contributed two data points for the upper and lower halves of the scratch. The shadows represent the actual upper and lower data points, while the central lines represent the average value of the well plate. (B) Cell scratch at 0 h. (C) Cell scratch at 54 h. Neon spherical cells indicate infected or dead cells directly captured by the light microscope. Quantification of the scratch area over time to model (D) cell migration and (G) infection rate in collagen alone (black) and collagen-coated liposomes (red). Likewise, (E) cell migration and (H) infection rates were measured under conditions with elastin alone (black) and elastin-coated liposomes (red). Similarly, (F) cellular migration and (I) infection development were evaluated for fibronectin alone (black) and fibronectin-coated liposomes (red).

Free collagen and collagen-coated liposomes drive comparable cell migration for roughly the first 30 hours. After this point, migration in the free collagen group slows and plateaus at 70% wound closure, whereas cells in the collagen-coated liposome group continue to migrate, ultimately reaching 89% coverage (Fig. 5D). In contrast, free elastin initially supports a migration rate similar to that of elastin-coated liposomes for the first 7 hours, then induces a faster migration over the following 21 hours before decelerating and ending at 82% coverage. Treatment with elastin-coated liposomes, however, enhances migration beginning around hour 28, resulting in 89% wound closure by the end of the assay (Fig. 5E). Analogous to the collagen results, free fibronectin and fibronectin-coated liposomes elicit similar migration rates during the first 15 hours. Thereafter, migration with free fibronectin slows and plateaus at 65% healing, while cells in the fibronectin-coated liposome group continue to migrate and achieve 83% closure by the end of the experiment (Fig. 5F). Relative to buffer-only and free liposome conditions, the ranking of final closure from most to least complete is as follows: elastin-coated liposomes (89%)*≃* collagen-coated liposomes (89%) *>* buffer only (85%) *>* fibronectin-coated liposomes (83%)*≃* free elastin (83%)*>* unmodified liposome (78.5%) *>* free collagen (70%) *>* free fibronectin (65%). These data underscore two key points: (1) free collagen and fibronectin do not interact sufficiently with cells to effectively enhance cell migration; and (2) liposomes coated with ECM proteins generally stimulate cell migration more effectively than either uncoated liposomes or the corresponding free proteins.

The pattern of bacterial infection rates varies with treatment conditions; nonetheless, the overall trend underscores liposomes’ role in limiting infection. In culture medium supplemented with free collagen, infection remains relatively low until approximately hour 30, then rises to a maximum of 0.6% and a minimum of 0.45%, before increasing again thereafter (Fig. 5G). When collagen-coated liposomes are applied, infection stays between 0.13–0.3% before climbing to 0.48% at the conclusion of the experiment. Treatment with free elastin leads to infection peaking at 0.8% at hour 37, then declining to 0.4% and subsequently rising again to 0.56% by the end of the assay (Fig. 5H). In contrast, elastin-coated liposomes maintain infection levels of 0.1–0.28%. Free fibronectin treatment results in contamination reaching 1.5% at the end of the assay, whereas fibronectin-coated liposomes keep contamination between 0.2–0.35% throughout (Fig. 5I). Relative to the two control conditions, the progression from lowest to highest infection at the end of the assays is: elastin-coated liposome (0.17%) *<* fibronectin-coated liposome (0.24%) *<* buffer alone (0.35%) *<* unmodified liposome (0.44%) *<* collagen-coated liposome (0.49%) *<* free elastin (0.56%) *<* free collagen (0.58%) *<* free fibronectin (1.5%). These findings indicate that liposomes themselves do not necessarily reduce infection, but rather serve as a platform on which ECM proteins, such as elastin and fibronectin, can unfold, making the membrane surface less permissive to infection. In contrast, ECM proteins in solution—particularly collagen and fibronectin—not only fail to interact effectively with cells and their secreted matrix to facilitate migration, but also actively increase infection.

## Discussion

### Individual Protein-Lipid Interactions

To develop an instructive scaffold environment, the ECM proteins employed are key to guiding cell behavior. Different mechanical properties and interactions of these proteins can be tailored to different mini-scaffold recipes to promote desired cell responses, whether that be growth and healing or cell differentiation. Prior to fabricating this pre-designed platform, the mechanoproperties of these proteins must first be characterized and understood to determine the most effective protein composition blends. The three proteins discussed in this study have been shown to possess different adsorption and mechanical behaviors and continue to differ in their binary and tertiary systems. The combined results from the interfacial elastic modulus, adsorption assays, X-ray reflectivity profiles, and cell scratch assays shed light on each protein’s unique role in the extracellular matrix and help to define their future application in mini-scaffolds.

Among these proteins, collagen is one of the most well-studied ECM proteins in terms of mechanical behavior and structure, and thus this characterization serves to validate and refine the collagen-lipid monolayer interaction model (Fig 6A). After spontaneous integration between the head and tail groups of the lipid, the start of compression showed a minor softening of the system (Fig. 1C). This protein, as discussed previously, significantly contributes to the strength and stability of the ECM, which collagen’s compression curve corroborates. This, however, appears to contradict the observed softening of the monolayer; the notable rise in elastic modulus at 35 mN/m surface pressure, however, supports literature findings. Collagen’s rigid rod structure allows the protein to possess high compressive stiffness and fracture resistance [36]. Instead, as compression begins, the proteins move and puncture through the monolayer, disrupting it, as indicated by increased roughness and decreased SLD (Table 1). The lipid monolayer eventually reassembles at 35 mN/m, at which point proteins can no longer rearrange and take up the compressive load imposed by the moving barriers (Fig. 1C). The residual sample at the bottom of the trough is most likely the lipid-protein sample that sank after disruption (Fig. S5A, Supplemental Information).

**Table 1.**
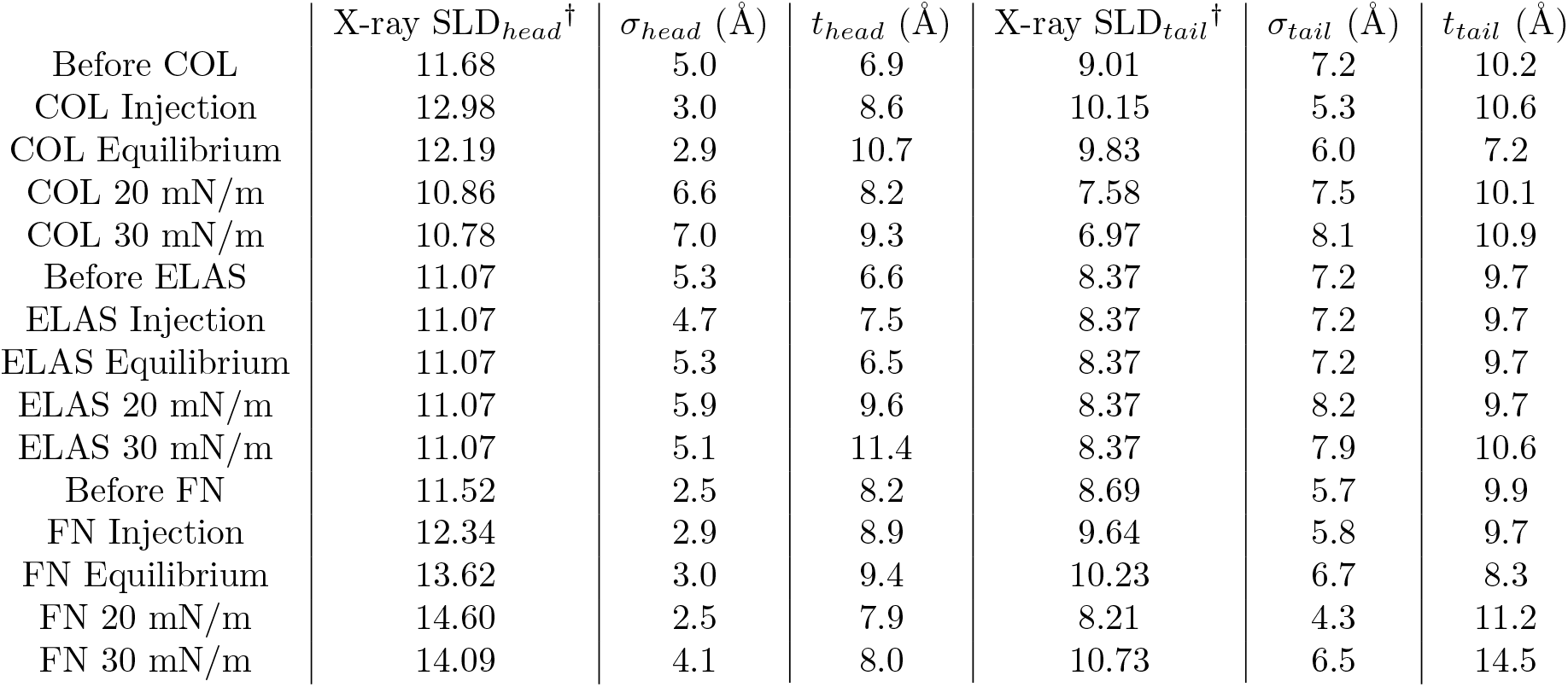
Summary of X-ray fitting parameters. ^†^ × 10^−6^/ Å ^2^.

**Figure 6.**
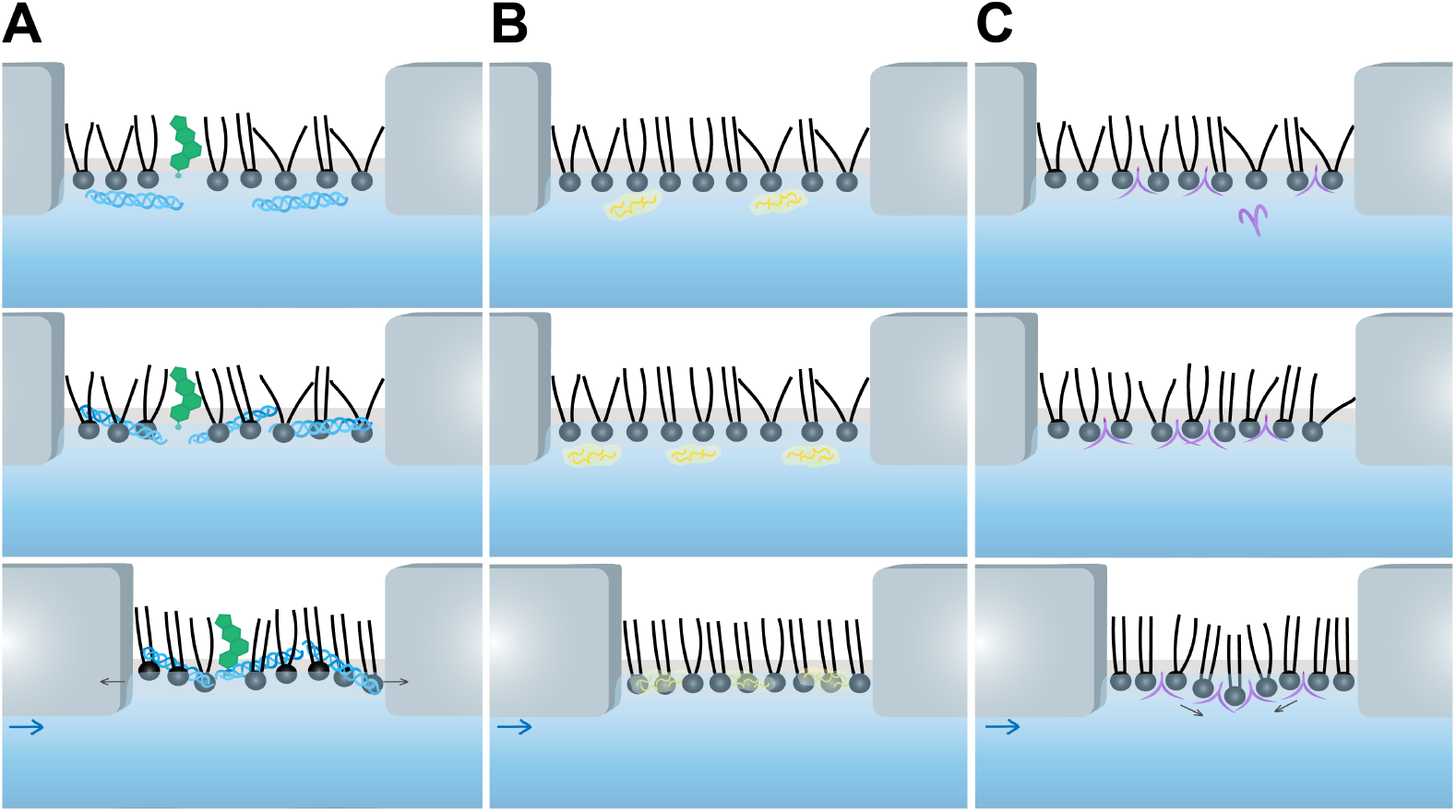
Schematic Representation of (A) collagen, (B) elastin, and (C) fibronectin proteins’ adsorption mechanisms in response to compressive forces. These models were determined using the adsorption assays and X-ray reflectivity profiles.

Elastin, whose primary recorded mechanical function is to store elastic energy, interacts much more passively with the lipid monolayer. The protein adsorbs more slowly than collagen and interacts with the head groups, only moving between the head and tail groups in response to compressive force, as evident by the changes in thickness and roughness of the headgroup layer (Fig 6B). Given the lack of residual protein in the trough between experiments, it is apparent that elastin is fully adsorbed onto the lipid film, allowing it to be aspirated along with the monolayer. This full incorporation of the protein with the lipid system is similar to that of collagen before compression, where collagen’s rigidity causes further disruption, but elastin’s mechanical properties don’t modify lipid film behavior while a compressive force is applied (Fig. 1D). The elastin-monolayer and pure monolayer systems collapse at the same pressure, demonstrating that the addition of elastin to the monolayer increases elastic modulus with no new mechanical behavior. This increase in modulus is due to an increase in the volume of material resisting compression, but the protein’s energy storage and elasticity cannot be observed during compression. After the collapse of the system, the elastic modulus drop is far higher than that of the lipid control, showcasing elastin’s “amplification” effect and energy storage as the protein permits greater folding and softens the membrane more.

As the only basement membrane protein out of the three studied, fibronectin showed a drastically different adsorption and interaction mechanism with the lipid monolayer (Fig 6C). At a slower rate than collagen (Fig. 1B), fibronectin adsorbs onto the lipid, settling between the head and tail groups, as indicated by dramatic increases in SLD of both head and tail layers (Table 1). In solution, fibronectin adopts a compact folded conformation, but upon adsorption it unfolds, enabling greater FN-FN binding [37]. Over time, the protein begins to form a network, weighing down the protein-bound monolayer and increasing the SLD and thicknesses of the head and tail group layers, as recorded in the x-ray reflectivity profiles (Fig. 4C and Table 1). Once compression begins in response to the mechanical stimulus, the protein network continues to assemble, dragging the monolayer downward. This result aligns with earlier reports showing that fibronectin can trigger vesicle budding upon binding to giant unilamellar vesicles [38]. As the film bulges inward, changes in roughness (Table 1) and the drop in elastic modulus (Fig. 1E) can be attributed to this mechanical interaction. The greater the compressive force, the greater the lipid film bulges, and eventually the lipid-protein bubbles drop to the bottom of the trough. This process is revealed by the first local minima in the compression curve (Fig. 1E) and in the protein-lipid remnants at the bottom of the trough (Fig. S5B, Supplemental Information). Breaks in the monolayer soften the system, and the monolayer stiffens only once the lipids reassemble into an uninterrupted film. For the rest of the compression, the remaining fibronectin network resists the compressive load applied to the system, stiffening it until a delayed collapse from the loss of material, suggesting fibronectin’s minor mechanical function in the ECM.

### Influences on Cellular Behaviors

The adsorption behavior and the structural and mechanical properties arising from protein–lipid interactions significantly influence the cellular responses observed in the scratch assay. Presentation of extracellular matrix (ECM) proteins on liposomes alters both migration dynamics and infection susceptibility compared to protein-only conditions. These findings suggest that not only protein identity, but also the mechanical and structural context in which proteins are presented, play a critical role in regulating cell behavior.

When introduced alone, these ECM proteins either exert minimal influence (elastin) or markedly inhibit cell migration (fibronectin and collagen). This is likely due to ineffective interactions with cells, which in turn diminish cellular motility. Notably, despite elastin and collagen possessing contrasting mechanical characteristics, their liposome-bound forms both enhance cell migration. The soft, elastic behavior of elastin at membrane interfaces aligns with the mechanical properties of skin cells, thereby facilitating fibroblast uptake during migration. Collagen on liposomes assembles in a more regulated manner [28], which improves fibroblast attachment during migration, whereas free collagen can spontaneously form large fibrils in solution, ultimately impeding cell movement. Treatment with fibronectin-coated liposomes slightly reduces the cell migration rate compared with the control. As a basement membrane protein, fibronectin exhibits a strong interaction with the lipid membrane and causes disruption (Fig. 4). When cells encounter fibronectin, its interaction with integrins can generate adhesion forces that can be excessively strong, thereby constraining cell motility [39].

The fluid and compliant nature of the lipid interface likely enabled dynamic ligand presentation, facilitating continuous adhesion turnover, which is necessary for efficient cell migration [40]. Enhanced migration in liposome-associated conditions may therefore result from optimized adhesion strength. Excessively strong adhesion can restrict movement, whereas insufficient adhesion limits traction generation [39]. The lipid-supported ECM proteins may provide an intermediate mechanical environment that promotes integrin engagement while maintaining adhesion plasticity, allowing cells to reorganize attachments and migrate more effectively across the wound area [41].

In addition to promoting migration, ECM proteins coated liposomes substantially reduce bacterial contamination across all three ECM systems. This observation is consistent with studies by Lee et al. and Chabria et al., where they similarly showed that stretched fibronectin can make the membrane surface less favorable for bacterial colonization, particularly Staphylococcus aureus (S. aureus), a frequent cause of opportunistic skin infections that can result in tissue damage [21, 42]. Another possible explanation is that altered protein conformation or spatial organization on lipid vesicles, which apply for all three ECM proteins, limits accessibility of viral binding sites or reduces receptor clustering required for efficient internalization [43]. Although our control group using unmodified liposomes did not exhibit a significant reduction in infection, earlier studies have demonstrated the antibacterial activity and infection-treating capabilities of liposomes [41, 44]. Together, these results demonstrate that protein–lipid interactions modulate both mechanical signaling and pathogen susceptibility in cultured cells. The ability to engineer ECM scaffolds through lipid platforms may provide a strategy to simultaneously enhance tissue regeneration while limiting infection risk, highlighting potential applications in wound healing, biomaterial coatings, and therapeutic delivery systems.

## Conclusions

In conclusion, our study systematically examined individual ECM proteins, examining how they structurally and mechanically affect lipid membranes and regulate cell migration. These proteins display distinct behaviors. Strong collagen adsorption on the membrane surface disrupts lipid packing, while its intrinsic rigidity yields an elastic modulus that is markedly higher than that of elastin or fibronectin. This stiff collagen network can form a robust scaffold for cell adhesion when its assembly is directed by the membrane, thereby enhancing cell attachment and promoting cell migration. In contrast, elastin exerts minimal structural or mechanical influence on the membrane during both adsorption and compression, yet it benefits cells by promoting migration and reducing infection. Fibronectin, on the other hand, exhibits complex mechanical responses to compression, associated with pronounced structural rearrangements initiated during adsorption. This strong interaction with the membrane, together with its specific binding to integrins on the cell surface, can generate excessively high adhesion forces that ultimately restrict cell motility.

The ratios of these proteins will later be tuned in the mixture to enhance cell growth, migration, and healing based on their mechanical behavior and interprotein interactions. Collagen is stiff in compression and provides the structural foundation for the ECM, while elastin, the other interstitial matrix protein, amplifies the mechanical properties of the ECM proteins by acting as a connecting molecule. Fibronectin, as a protein that interacts with both cell receptors and interstitial proteins, forms large networks and is more responsive to external stresses, but it cannot provide mechanical stability. These findings provide a better understanding of how the ECM interacts mechanically with itself, enabling us to begin manipulating and hastening ECM secretion. Additionally, knowledge of these proteins’ interactions and additive capabilities can enable the production of a strategically designed native scaffold and various materials tailored for specific biomedical applications.

## Author Contributions

M.D.P. designed the research. V.I. carried out Langmuir isotherms, protein adsorption assays, and performed interfacial elastic calculations. V.I. and J.R. performed and analyzed scratch assays. V.I. and M.D.P. carried out X-ray reflectivity experiments. M.D.P. analyzed and modeled X-ray data. V.I, J.R., and M.D.P wrote the article.

## Acknowledgments

Support for M.D.P. was provided through the Midscale RI:1 program of the NSF, Award DMR-1935956. Support for V.I. was provided by the Center for High Resolution Neutron Scattering, a partnership between the National Institute of Standards and Technology and the National Science Foundation under Agreement No. DMR-2010792. This research was also partially supported by funds from the University of Missouri Research Reactor through the Nuclear Science Career Academy Summer 2025 Internship Program. We thank Dr. Sushil Satija for the help in setting up the X-ray instrument at the NIST Center for Neutron Research (NCNR). Additionally, V.I. thanks Julie Borchers, Donna Kalteyer, and Juscelino Leao for their support during the SURF 2024 Internship program at NCNR. We are deeply grateful for the assistance in cell experiments from Dr. Taixing Cui’s lab at Mizzou’s Dalton Cardiovascular Research Center, particularly Dr. Yixiao Liu and Dr. Jianxin Yan. The commercial products used in this study were referenced only to adequately specify the experimental procedure. Such identification of commercial products is not intended to imply recommendation or endorsement by the National Institute of Standards and Technology, nor is it intended to imply that the identified products are necessarily the best available for the purpose. A preliminary version of this work, [DOI: 10.64898/2026.04.03.716427], was deposited in bioRxiv on 4/6/2026.

