## Supplemental Information for "Binding Structures, Mechanical Properties, and Effects on Cellular Behaviors of Extracellular Matrix Proteins on Biomembranes"

**S1**. Scratch Assay Migration Analysis on Image J

All scratch assay images were processed for migration analysis using ImageJ. The scale was set to 1296 pixels per 1000 μm, as indicated by the scale bar at the bottom-right corner of each photo. Using the polygon area selection tool, the cell scratch boundary was traced manually, and the measurement tool was used to calculate the area. All areas were normalized. The Mean was calculated by averaging the upper and lower images of the same well and graphed against time. The actual area values of the upper and lower images are used to produce the error.

$$\text{Normalization: } \frac{A_t}{A_0} \times 100$$

$$\text{Mean: } \frac{Upper + Lower}{2}$$

$$\text{Standard Deviation: } \sqrt{\frac{(Upper - Mean)^2 + (Lower - Mean)^2}{2}}$$

Where $A_t$ is the area of the hour of interest, $A_0$ is the initial area (hour 3), Upper is the normalization of the area above the drawn center line, and Lower is the normalization of the area below the drawn center line.

**S2.** MATLAB Code

MATLAB code was developed to determine the number of infected cells in each scratch image. All files are loaded in from the designated folder and are ordered by hour. The code was designed to isolate infected cells (which appear neon yellow/green in color) with a target [R G B] of [195, 255, 0] and divided by 255 to normalize the value between 0 and 1. The tolerance of this color was set to 0.3. A grayscale image or a mask for each file was created to extract only the photo's pixels within the color range. Using the image's pixel-to-area scale, the area of infected cells can be measured, and dividing by the total number of pixels yields the percentage of infected cells within the image. Each mask image was saved, and diagnostics were printed in Excel.

### **S3**. Full Scratch assay Images

Tris-HCl Buffer condition:

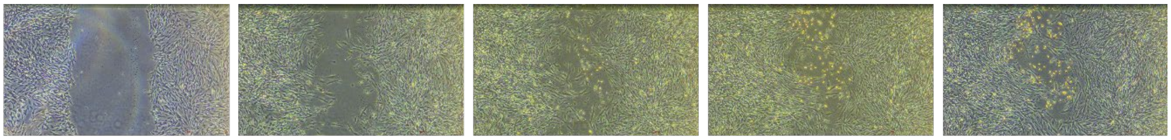

Hour 3 Hour 18 Hour 30 Hour 42 Hour 54

Uncoated liposome condition:

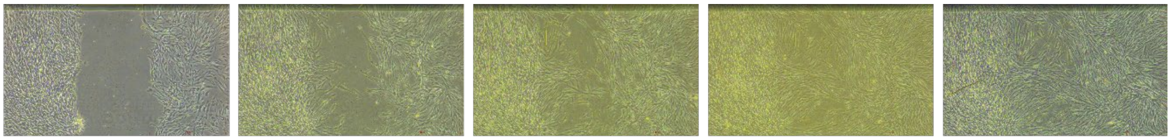

Hour 3 Hour 18 Hour 30 Hour 42 Hour 54

Free collagen condition:

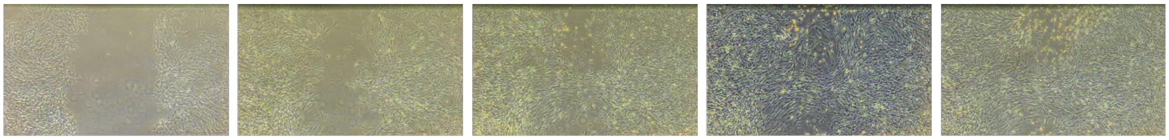

Hour 3 Hour 18 Hour 30 Hour 42 Hour 54

Collagen-coated liposome condition:

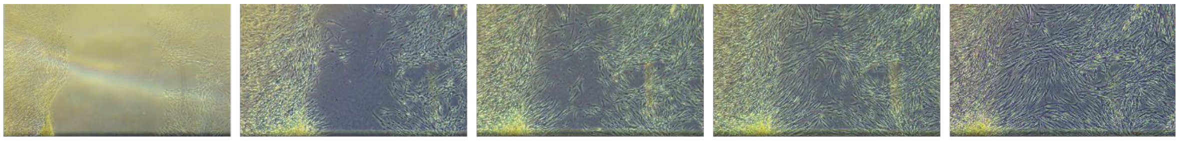

Hour 3 Hour 18 Hour 30 Hour 42 Hour 54

Free elastin condition:

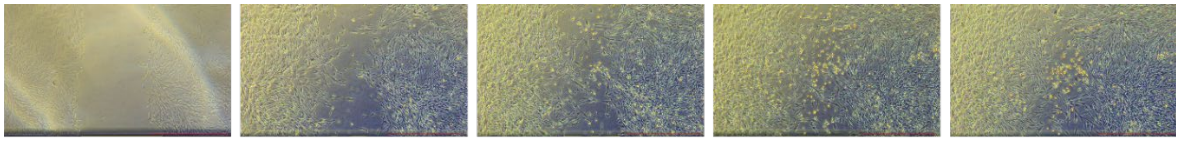

Hour 3 Hour 18 Hour 30 Hour 42 Hour 54

Elastin-coated liposome condition:

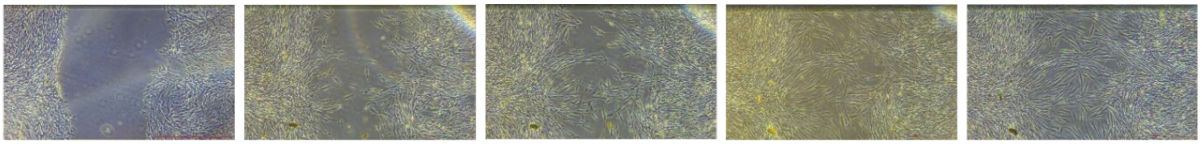

Hour 3 Hour 18 Hour 30 Hour 42 Hour 54

Free fibronectin condition:

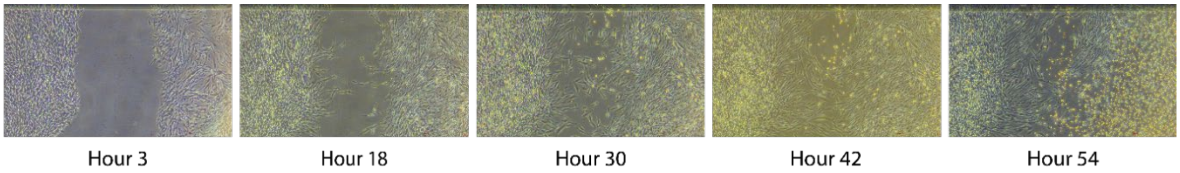

Fibronectin-coated liposome condition:

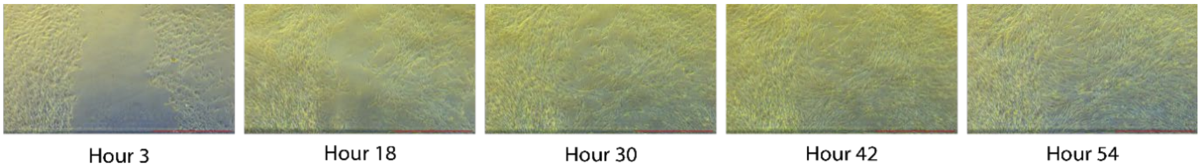

Spherical neon cells indicate infected or dead cells. The change in image color results from changes in pH. Dead/infected cells cause acidity, lowering pH in culture, resulting in a brighter/yellow hue to images.

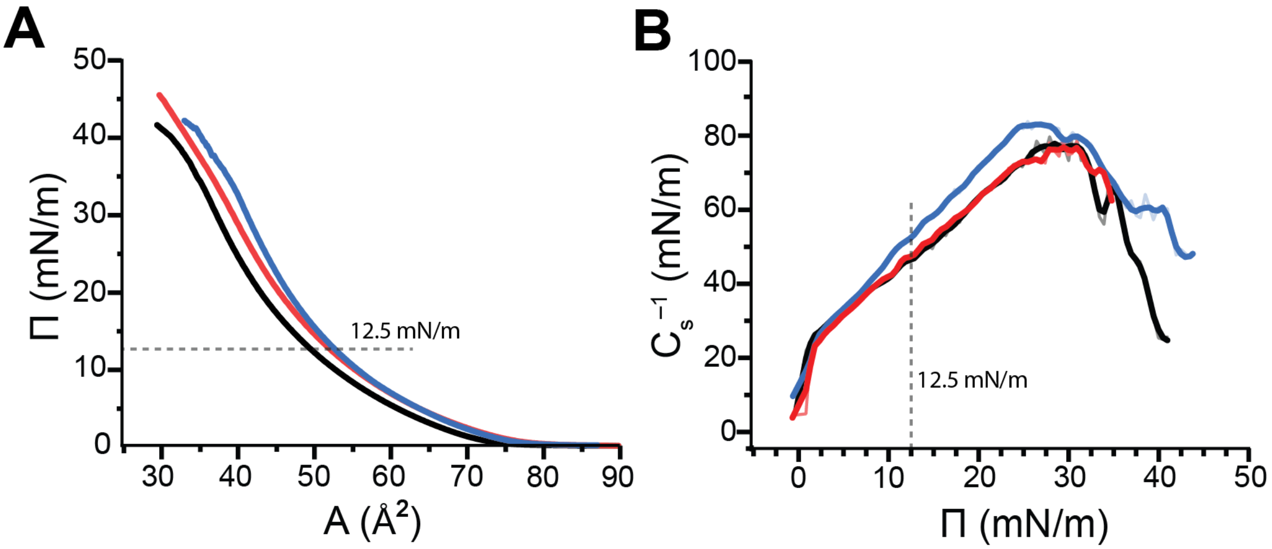

**Figure S4.** Surface pressure-area isotherms (Π-A isotherms) of various DMPC mixed monolayers at an air-water interface. The black curve is DMPC; the red curve is DMPC+DMPS; and the blue curve is DMPC+DMPS+Chol. (B) Surface pressure Π versus the elastic moduli $C^{-1}$ of area compressibility calculated from Π-A isotherms for DMPC mixed monolayers. The dashed line, for visual purposes, indicates the chosen pressure for protein deposition.

### Langmuir Isotherms Characterization

The liftoff area, the molecular area where the isotherms deviate from their baseline, indicates the onset of intermolecular interactions, and is approximately the same for DMPC, DMPC with DMPS (DMPC+DMPS), and DMPC with DMPS and cholesterol (DMPC+DMPS+Chol). There are no plateaus on these isotherms, suggesting that the films do not undergo a liquid-expanded-to-liquid-condensed phase transition but rather remain in a liquid-expanded state until the collapse pressure is reached. This phase behavior is well observed for PC lipids with short acyl chains (1). Compared to DMPC, the two protein-binding platforms (DMPC+DMPS and DMPC+DMPS+Chol) packed less effectively and had similar packing densities up to a surface pressure of 12.5 mN/m, which was chosen for ECM protein deposition. DMPC, DMPC+DMPS, and DMPC+DMPS+Chol. took off at $75.6 \pm 2.4$, $78.0 \pm 1.5$, and $76.5 \pm 2.5$ Å$^2$ in the molecular area, respectively. Despite a negligible difference in packing density, the mechanical behaviors were noticeably different at 12 mN/m as shown in Figure S4. The elastic modulus of the cholesterol-containing lipid mixture is higher than that of the cholesterol-free lipid mixture, consistent with previous findings (2). The elastic modulus curves also show a similar trend: the lipid mixture containing cholesterol has an overall higher modulus and stiffness than the other two curves, supporting findings from previous studies (2). The peak elastic moduli for all three lipid systems are at approximately 30 mN/m, and are representative of a biological bilayer system (3, 4). The similarity in the curvature of the lipid models confirms the reproducibility of Langmuir isotherms and enables effective comparison of protein-lipid interactions in lipid systems with and without cholesterol, while accounting for changes in stiffness. These results provide a foundational basis for analyzing protein-lipid models and characterizing their mechanical properties.

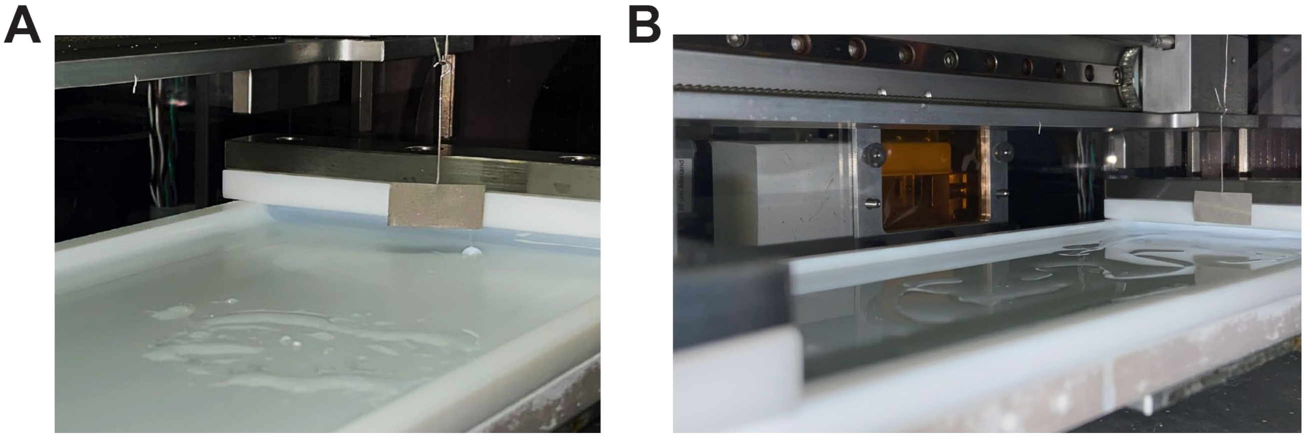

**Figure S5**. Images depict the usage of the Langmuir Trough to describe the surface pressure (Π) vs. Time adsorption assay. Individual proteins were injected beneath the film and allowed to adsorb for 24–72 hours. After removing the buffer, the residue of the lipid monolayer film can be observed on the trough's surface. These indicate material that failed to remain on the liquid surface and was not removed by simple aspiration. Residues on the trough surface after (A) collagen and (B) fibronectin adsorption experiments.
